## Supplementary information for "Death Induced by Survival gene Elimination (DISE) is correlated with neurotoxicity in Alzheimer’s disease and aging"

Brief description of what this file includes:

Supplementary figure legends

Supplementary Fig. 1 | SPOROS analysis of total sRNAs in two AD mouse models.

Supplementary Fig. 2 | Total read numbers of neuron, glia and immune cell specific RISC bound miRNAs in different brain samples.

Supplementary Fig. 3 | Repeat analysis of young and old mouse brains using SPOROS.

Supplementary Fig. 4 | Characterization of human iPSC-derived midbrain dopamine neurons that were aged *in vitro*.

Supplementary Fig. 5 | 6mer seed viability of R-sRNAs in three normal and three SuperAger brains.

Supplementary Fig. 6 | Differentiated SH cells are more sensitive to the toxicity of Aβ42.

Supplementary Fig. 7 | SPOROS analysis of total sRNAs in Drosha k.o. NB7 cells.

Supplementary Fig. 8 | ATA blocks RISC uptake of DISE-inducing sRNAs and protects from DISE in NB7 and SH cells.

Supplementary Fig. 9 | Characterization of Ago2 k.o. SH cells.

Supplementary Fig. 10 | All uncropped Western blots.

Supplementary Table 1 | Description of human brain material used.

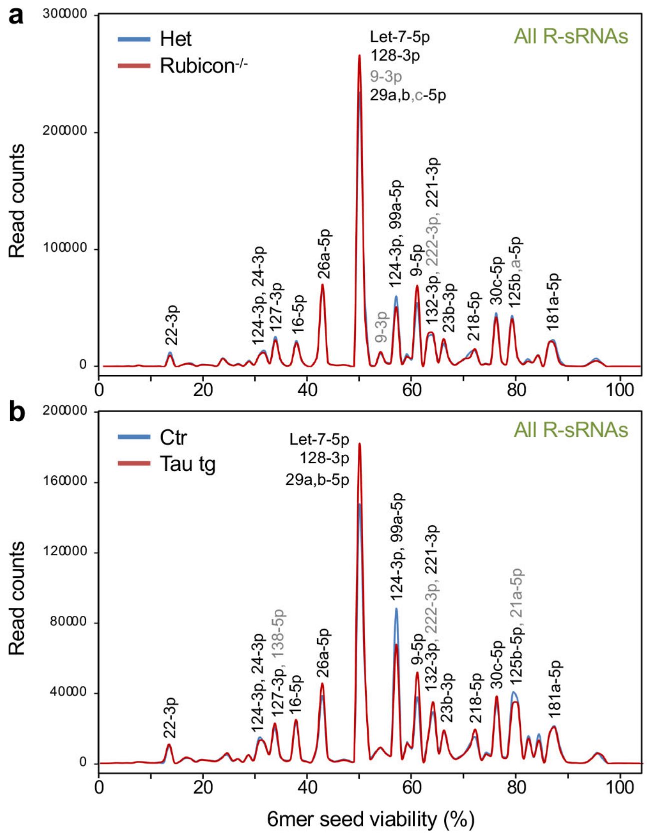

**Supplementary Fig. 1. SPOROS analysis of total sRNAs in two AD mouse models. a, b,** 6mer seed viability graph of total R-sRNAs of 5XFAD Rubicon$^{+/-}$ (Het) and 5XFAD Rubicon$^{-/-}$ (**a**) and Ctr and Tau transgenic (tg) (**b**) brains. Shown are averages of triplicate (**a**) and quadruplicate (**b**) samples. In all cases sRNAs with 10,000 or more reads in at least one sample are labeled. In each label sRNAs are listed in the order of expression levels. sRNAs shared in both analyses are shown in black.

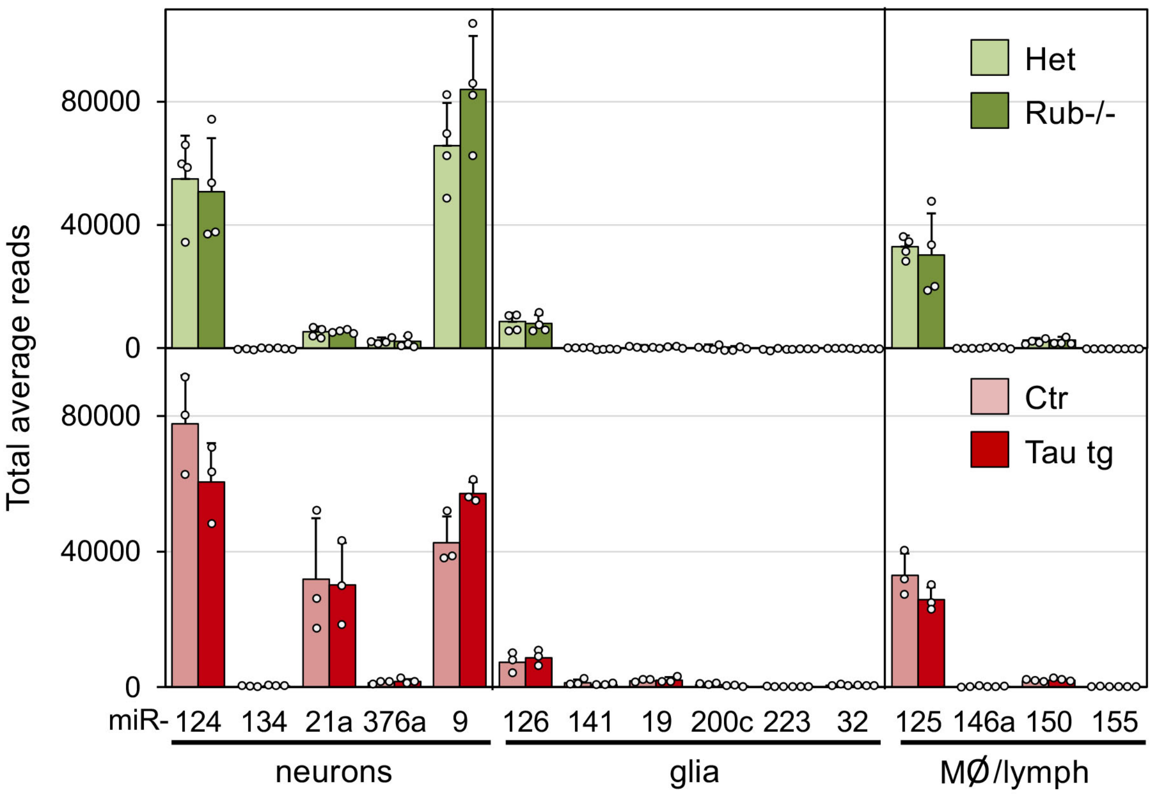

**Supplementary Fig. 2. Total read numbers of neuron, glia and immune cell specific RISC bound miRNAs in different brain samples.** Normalized expression levels of miRNAs pulled down in different tissues in the two experiments described in Fig. 1d, e. Shown are averages with SD. None of the comparisons between control and respective AD model reached statistical significance.

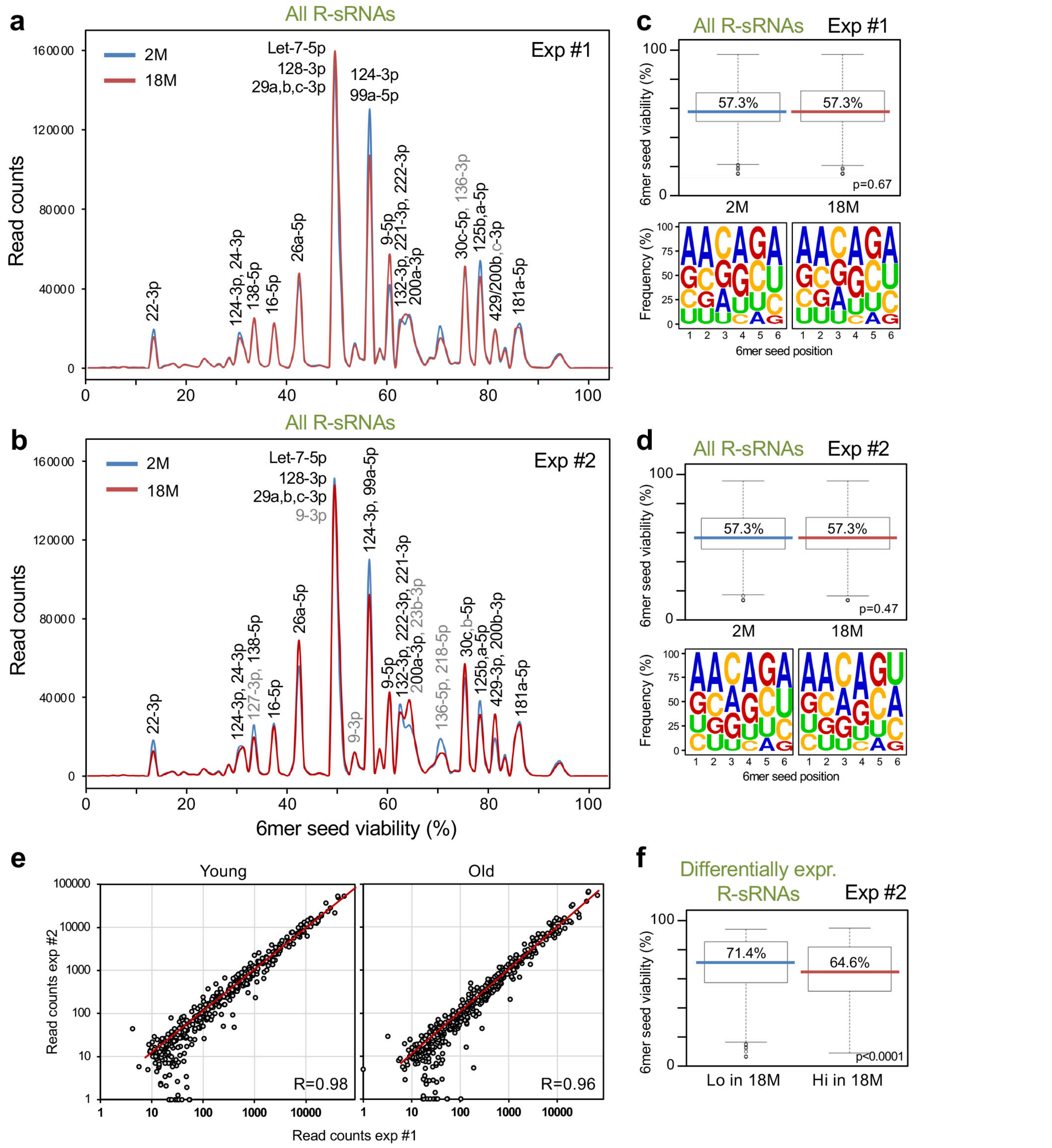

**Supplementary Fig. 3. Repeat analysis of young and old mouse brains using SPOROS. a, b,** 6mer seed viability graph of total R-sRNAs of 2-month and 18-month old mouse brains. Experiment #1 (**a**) and Experiment #2 (**b**). Shown are averages of triplicate (**a**) and duplicate (**b**) samples. In all cases sRNAs with 10,000 or more reads in at least one sample are labeled. In each label sRNAs are listed in the order of expression levels. sRNAs shared in the two analyses are shown in black. **c, d,** *Top*, 6mer seed viability plots of the total 6mer seed viability of the samples in **a** and **b**. Medians (in the color of each sample/condition) and Kruskal-Wallis median test p-value are given. *Bottom*, average 6mer seed composition of the sRNAs in the same samples. **e,** Pearson correlation analysis of gene products covered by the reads precipitated in the experiments shown in **a** (Exp #1) and in **b** (Exp #2). Only genes were included with an average of at least 10 reads across all samples. **f,** 6mer seed viability plot of differentially expressed R-sRNAs of 2-month and 18-month old mouse brains (Exp. #2). Medians (in the color of each sample/condition) and Kruskal-Wallis median test p-value are given. Definition of box and whisker plots in c, d, f: The lower and upper hinges of the box correspond to the first and third quartiles (the 25th and 75th percentiles). The upper/lower whiskers extend from the upper/lower hinge to the largest/smallest value no further than 1.5 x inter-quartile range (distance between the first and third quartiles) from the hinge. Data beyond the end of the whiskers are outliers and are plotted individually.

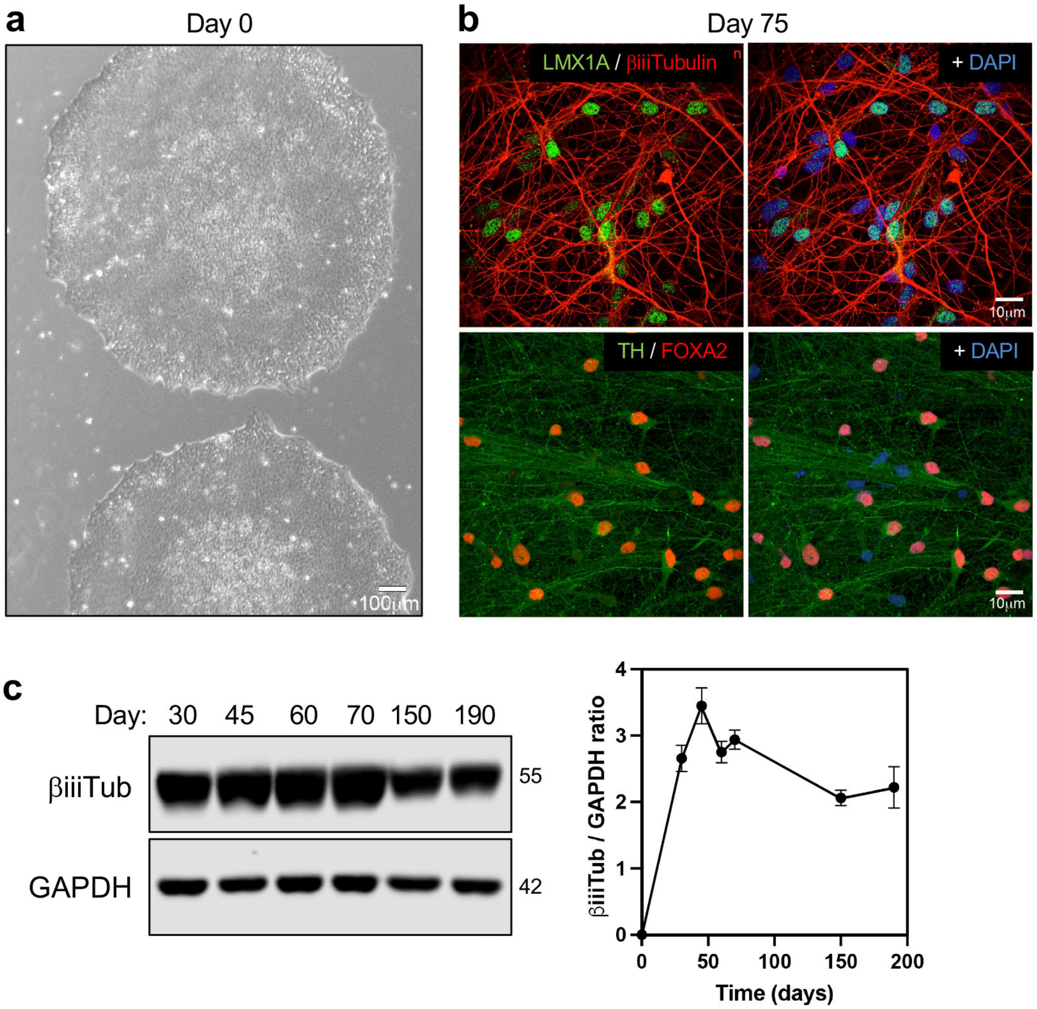

**Supplementary Fig. 4. Characterization of human iPSC-derived midbrain dopamine neurons that were aged *in vitro*. a,** Brightfield image of iPSCs before differentiation (day 0). **b,** Immunostaining analysis of midbrain dopamine (DA) neurons at day 75 using antibodies that detect neuronal (βiii Tubulin) or DAergic specific markers (LMX1A, FOXA2, Tyrosine Hydroxylase (TH). DAPI (blue) was used to stain nuclei. **c,** *Left,* Western blot analysis of cultured neurons for neuronal marker βiiiTubulin and loading control, GAPDH. *Right*, Quantification of western blot.

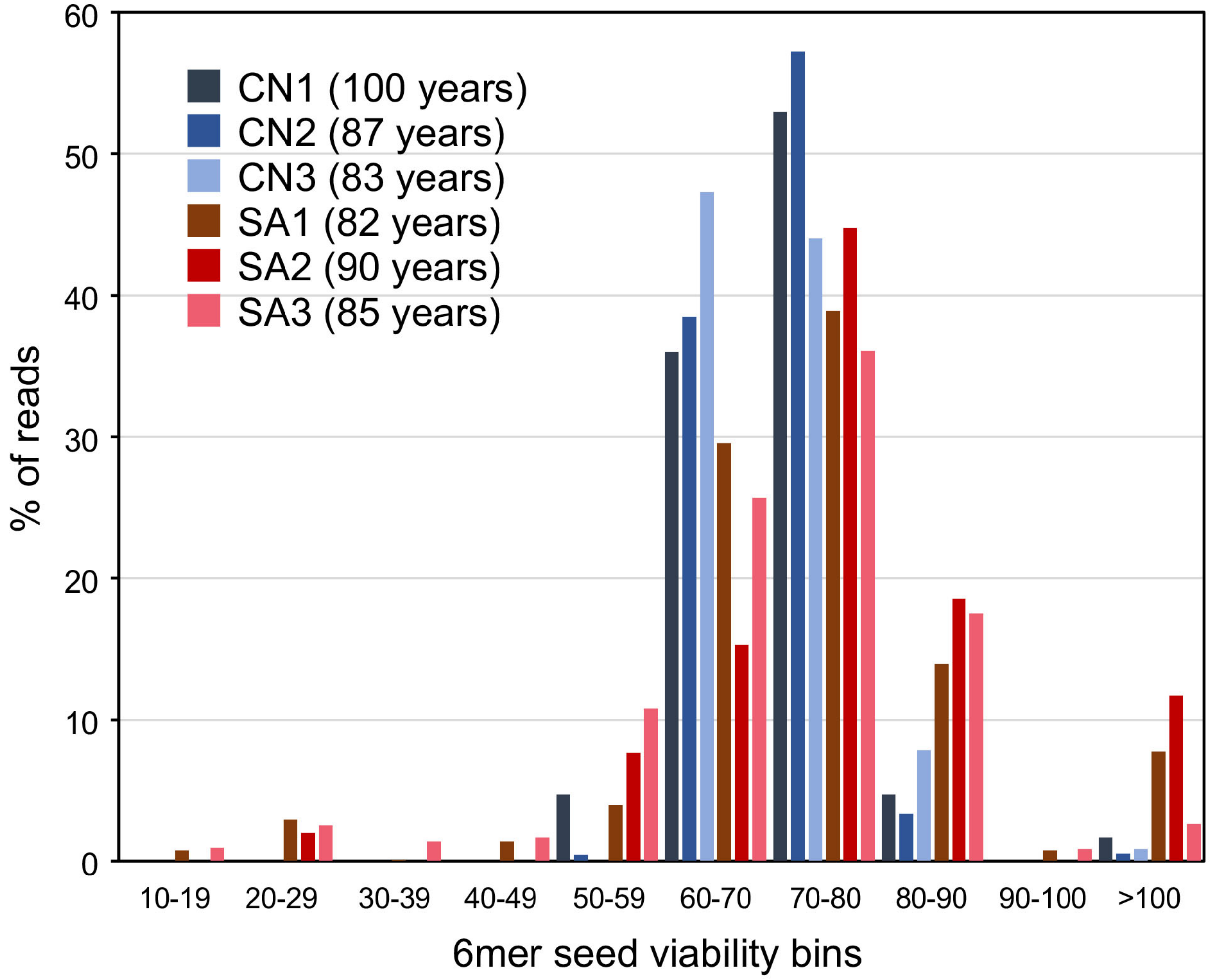

**Supplementary Fig. 5. 6mer seed viability of R-sRNAs in three normal and three SuperAger brains.** Breakdown of the data in Fig. 3d on R-sRNAs differentially expressed between 3 cognitive normal (CN) and 3 SuperAger (SA) brains. 6mer seed viabilities are shown as percent of total reads and sorted into bins of 10% 6mer seed viability. The age of each participant is shown in brackets. For details on participants see Supplementary Table 1.

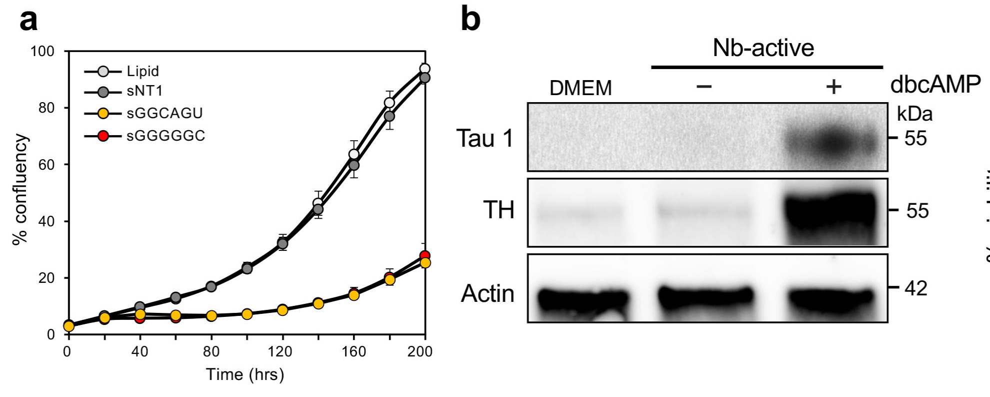

**Supplementary Fig. 6. Differentiated SH cells are more sensitive to the toxicity of Aβ42. a,** Change in confluence over time of undifferentiated SH cells treated with lipid only or transfected with 10 nM of the indicated sRNA. Data points are from quadruplicate and mean -/+ SE. This is representative of three independent experiments. **b,** Western blot analysis of SH cells cultured in DMEM or in NB-active medium with and without dbcAMP. TH, tyrosine hydroxylase. **c,** Viability assay of dbcAMP day 1 or day 7 differentiated SH cells left untreated or treated with either Aβ40 or Aβ42 for 48 hrs. Data are in triplicate. This is representative of two independent experiments. Shown is the mean with SD and Student's t-test p-values.

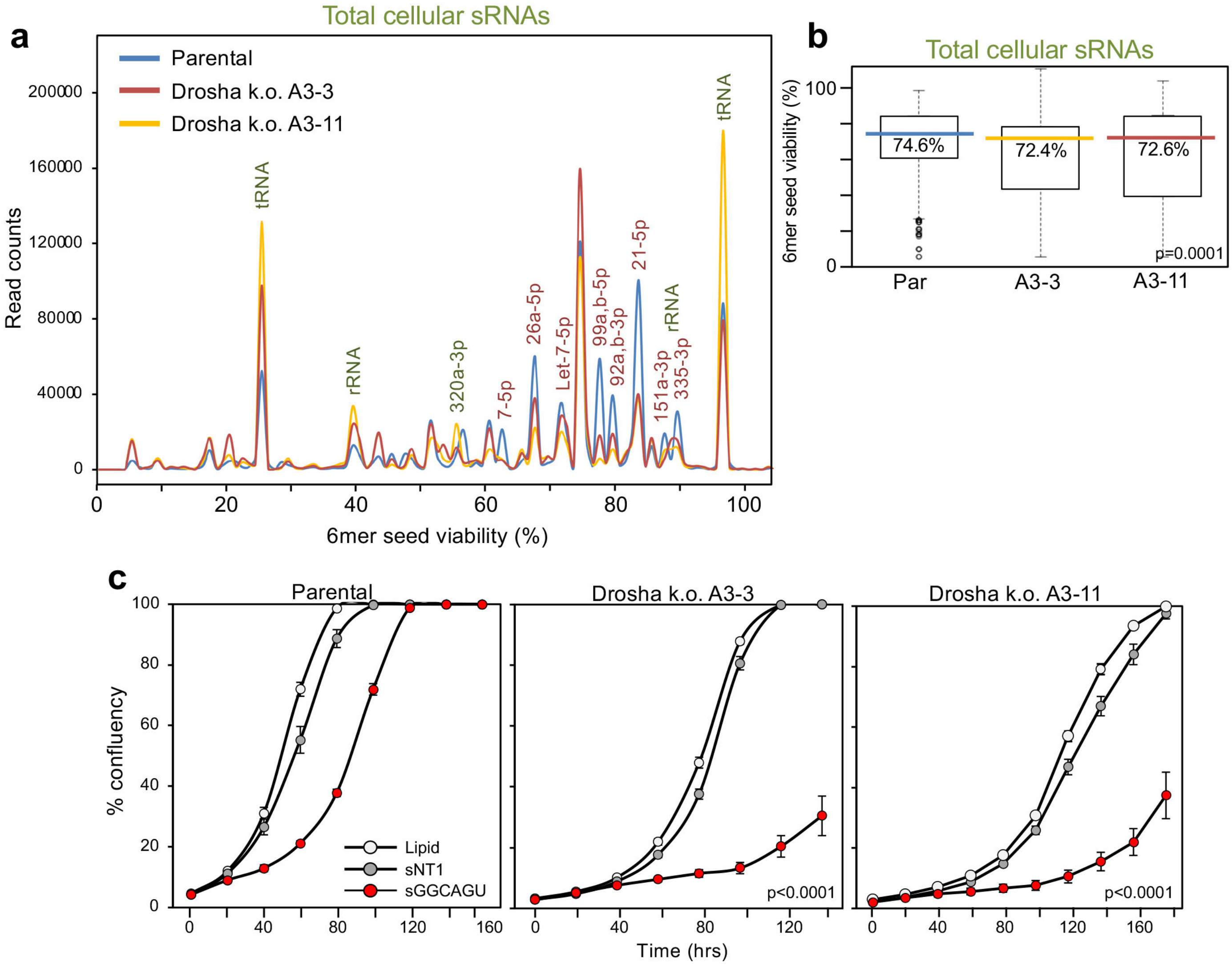

**Supplementary Fig. 7. SPOROS analysis of total sRNAs in Drosha k.o. NB7 cells. a,** 6mer seed viability graph of all sRNAs sequenced in parental NB7 cells and two Drosha k.o. clones. sRNAs with 10,000 or more reads in at least one sample are labeled. Enriched sRNAs are shown in green and depleted sRNAs in red. **b,** 6mer seed viability plots of the 6mer seed viability of the samples in **a**. Shown are the medians (in the color of each sample/condition) and Kruskal-Wallis median test p-value. For definition of box and whisker plot see legend of Supplementary Figure 3. **c,** Change in confluence over time of NB7 parental cells or two Drosha k.o. clones treated with lipid only or transfected with 1 nM of siGGCAGU. Data points are from triplicates and mean -/+ SE. P-values were calculated using a polynomial distribution test and support a difference in sensitivity between parental NB7 and the two Drosha k.o. clones.

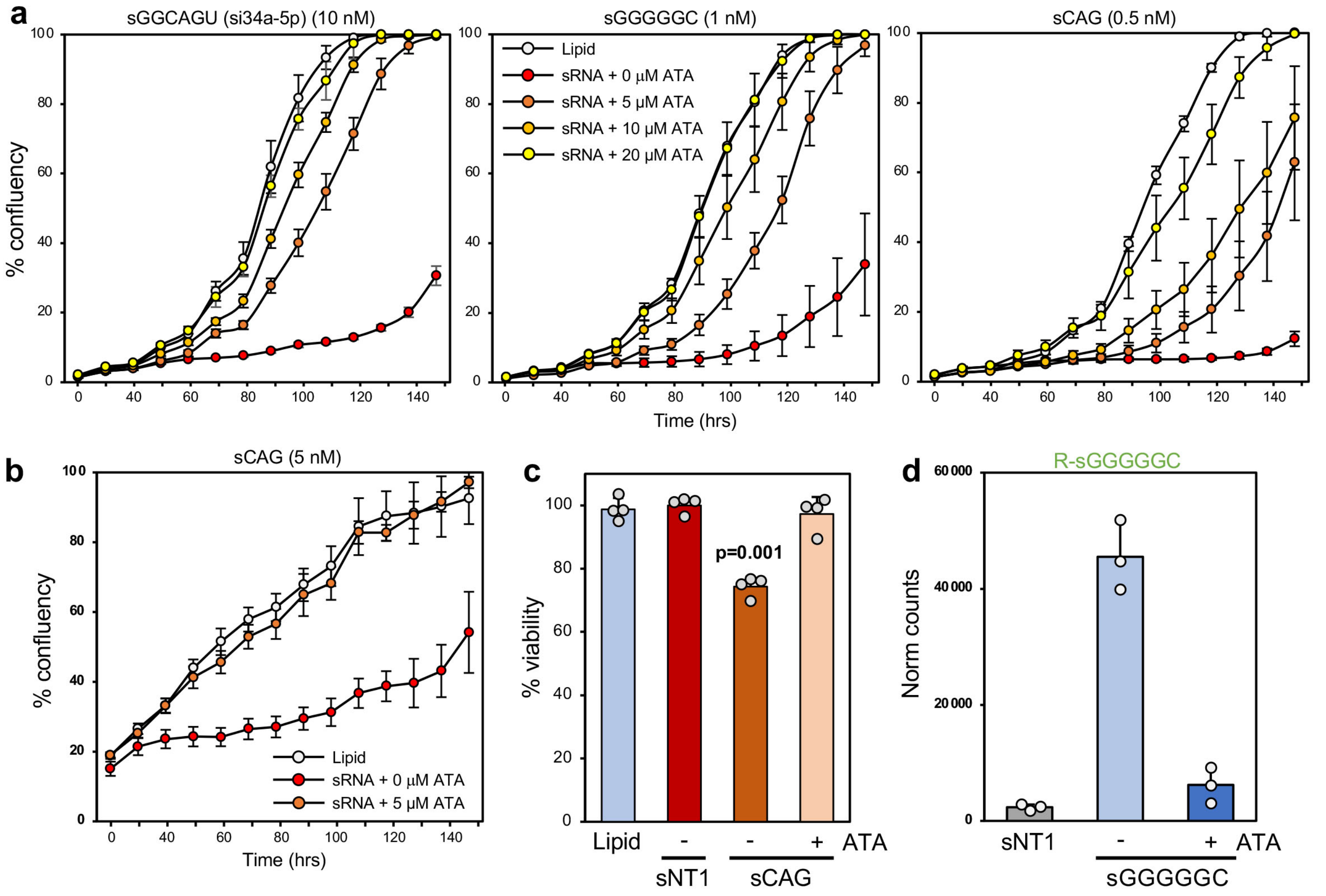

**Supplementary Fig. 8. ATA blocks RISC uptake of DISE-inducing sRNAs and protects from DISE in NB7 and SH cells. a,** Change in confluence over time of NB7 cells treated with lipid only or transfected with the indicated sRNA in the absence or presence of increasing concentrations of aurintricarboxylic acid (ATA). Data points are from triplicates and mean -/+ SE. Data are representative of three independent experiments. **b,** Change in confluence over time of SH cells treated with lipid only or transfected with 5 nM of sCAG in the absence or presence of 5 μM ATA. Data points are from triplicates and mean -/+ SE. **c,** Viability assay of differentiated SH cells 72 hrs after treatment with either lipid or 5 nM of sNT1 or sCAG in the absence of presence of 10 μM ATA. Student's t-test p-value is shown for the comparison between cells treated with sNT1 versus sCAG. **d,** Counts (normalized to 1 million reads) of the sGGGGGC antisense strand in the RISC of NB7 cells transfected with either 1 nM sNT1, sGGGGGC or sGGGGGC and treated with 10 μM ATA. Shown are the averages of biological triplicates -/+ SD.

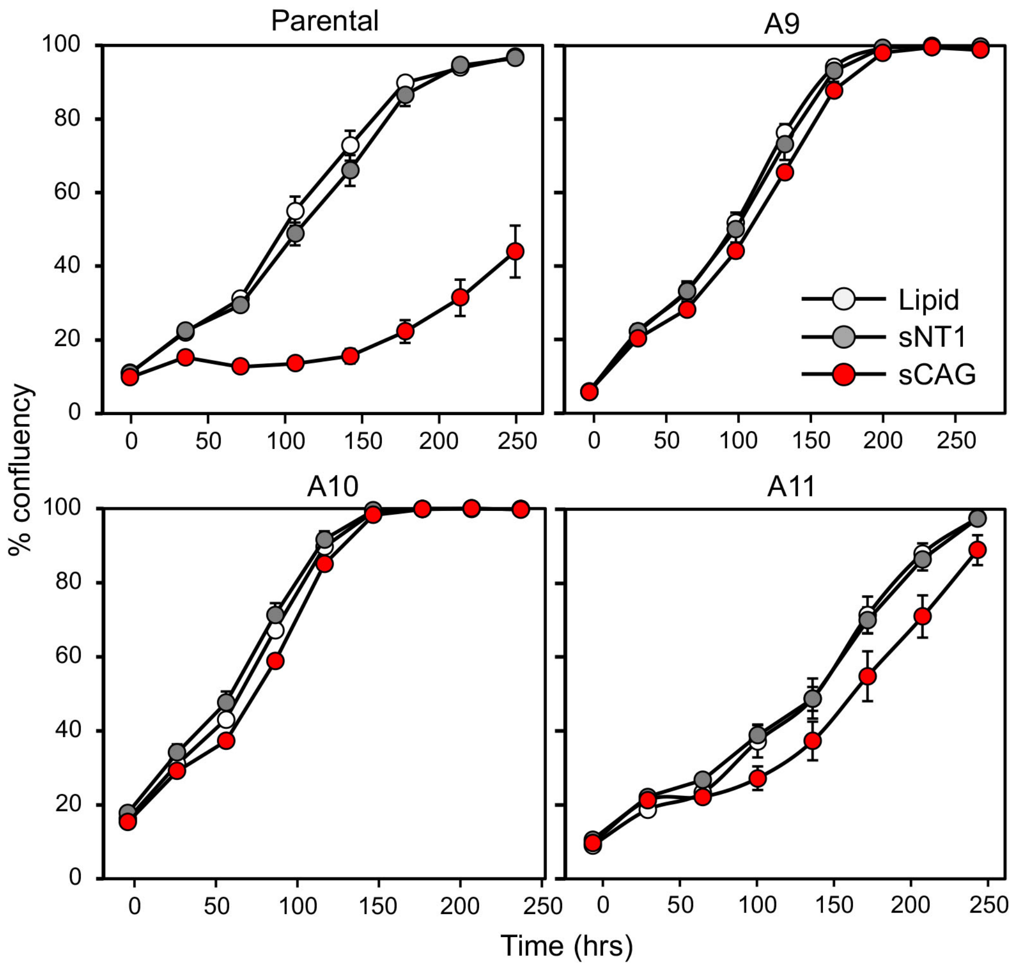

**Supplementary Fig. 9. Characterization of Ago2 k.o. SH cells.** Change in confluence over time of SH parental and Ago2 k.o. cells treated with lipid only or transfected with 5 nM of the indicated sRNAs. Data points are from triplicates and mean -/+ SE. Data were confirmed in another independent k.o. clone.

Fig. 2c

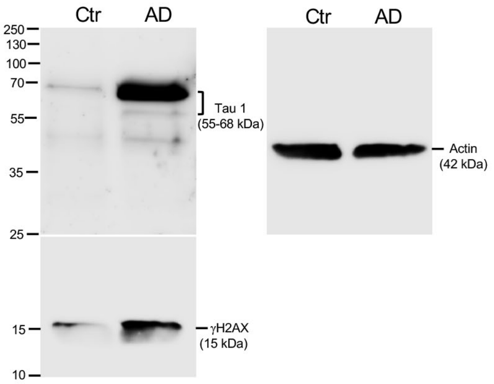

Fig. 4b

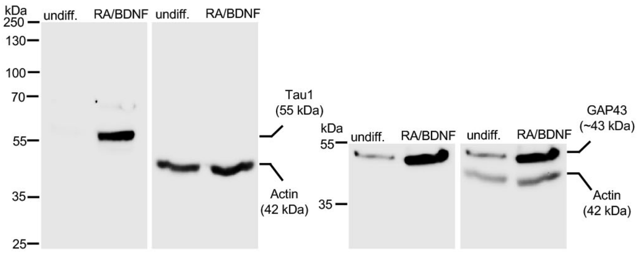

Fig. 3b

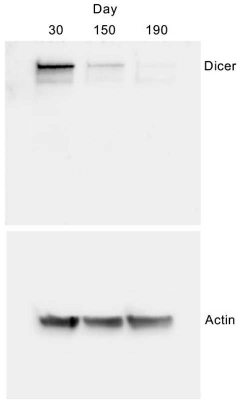

Fig. 5a

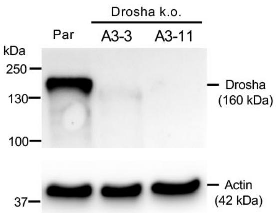

Fig. 6b

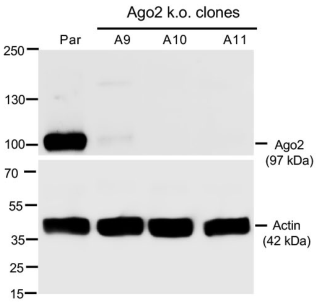

Fig. 6c

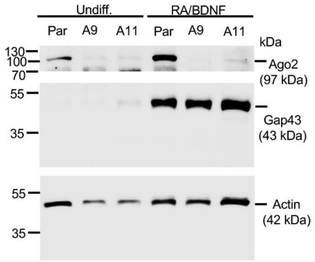

Fig. 6e

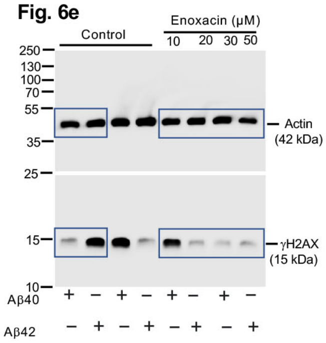

Fig. 6h

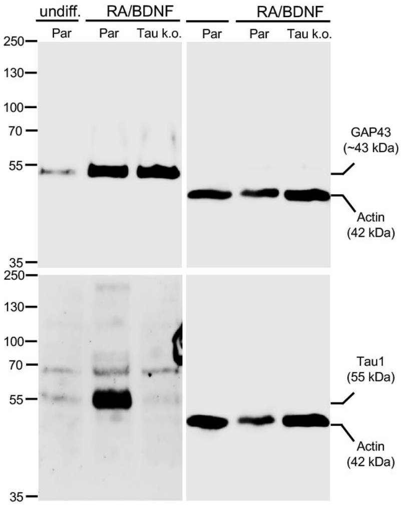

Fig. 6f

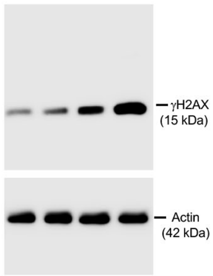

Fig. 6g

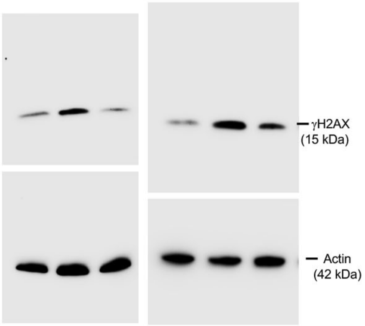

Fig.6j

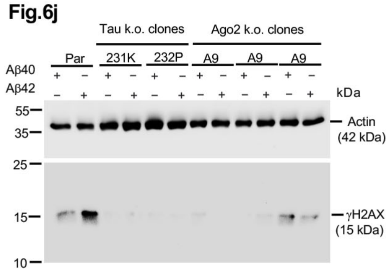

Fig. 6k

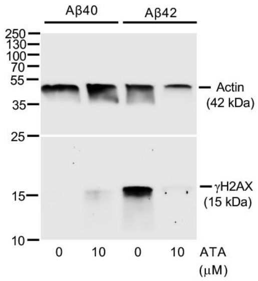

Fig. S4c

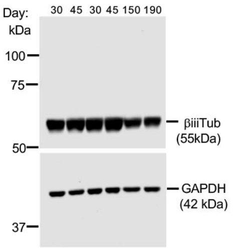

Fig. S6b

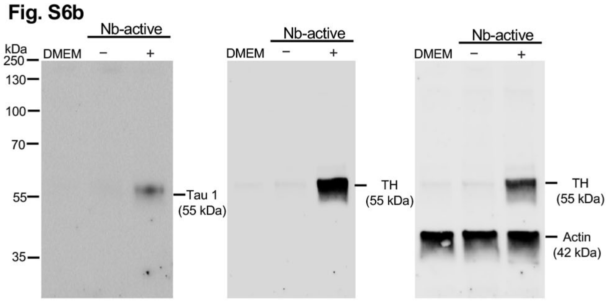

**Supplementary Fig. 10. All uncropped Western blots.**

| | ID | Region | ApoE | Age | Gender | PMI | Race | Clinical Diagnosis | Br wt | Pathologic Diagnosis |
|---|---|---|---|---|---|---|---|---|---|---|
| **Controls** | | | | | | | | | | |
| | CN1 | T | | 83 | Fe | 43 | C | CN | 1240Fr | Low ADNC |
| | CN2 | T & F | | 87 | M | 16 | C | CN | 1400Fr | Intermediate ADNC |
| | CN3 | T & F | | 100 | Fe | 14 | C | CN | 1000Fr | Intermediate ADNC |
| Average | | | | 90 | | 24.3 | | | | |
| **SuperAgers** | | **Region** | | **Age** | **Gender** | **PMI** | **Race** | **Clinical Dx** | **Weight** | **Pathologic Diagnosis-1** |
| | SA1 | F | | 90 | M | 14 | C | CN (SA) | 1267Fr | Low ADNC |
| | SA2 | F | | 85 | Fe | 9 | C | CN (SA) | 1310Fr | Low ADNC Tau Only |
| | SA3 | F | | 82 | Fe | 24 | C | CN (SA) | 1241Fr | Low ADNC |
| Average | | | | 85.7 | | 15.7 | | | | |

**Supplementary Table 1**. Description of human brain material used. CN = Cognitive normal, SA = SuperAger, T = Temporal Cortex, F = Frontal lobe, M = Male, Fe = Female, PMI = Postmortem Interval, C = Caucasian, Fr = Fresh, ADNC = Alzheimer's Disease Neuropathologic Change.
